# Patient-derived glioblastoma organoids as real-time avatars for assessing responses to clinical CAR-T cell therapy

**DOI:** 10.1101/2024.10.03.616503

**Authors:** Meghan Logun, Xin Wang, Yusha Sun, Stephen J. Bagley, Nannan Li, Arati Desai, Daniel Y. Zhang, MacLean P. Nasrallah, Emily Ling-Lin Pai, Bike Su Oner, Gabriela Plesa, Donald Siegel, Zev A. Binder, Guo-li Ming, Hongjun Song, Donald M. O’Rourke

## Abstract

Patient-derived tumor organoids have been leveraged for disease modeling and preclinical studies, but rarely applied in real-time to aid with interpretation of patient treatment responses in clinics. We recently demonstrated early efficacy signals in a first-in-human, phase 1 study of dual-targeting chimeric antigen receptor T cells (EGFR-IL13Rα2 CAR-T cells) in patients with recurrent glioblastoma. Here we analyzed six sets of patient-derived glioblastoma organoids (GBOs) treated concurrently with the same autologous CAR-T cell products as patients in our phase 1 study. We found that CAR-T cell treatment led to target antigen reduction and cytolysis of tumor cells in GBOs, the degree of which correlated with CAR-T cell engraftment detected in patients’ cerebrospinal fluid (CSF). Furthermore, cytokine release patterns in GBOs mirrored patient CSF samples over time. Our findings highlight a unique trial design and GBOs as a valuable platform for real-time assessment of CAR-T cell bioactivity and insights into immunotherapy efficacy.

## INTRODUCTION

Patient tumor-derived organoids have been increasingly recognized as an essential tool for cancer research, providing a more accurate and physiological representation of patient tumors compared to cell line or xenograft models^1–3^. While tumor organoids have been leveraged to study biological mechanisms of cancer^4^ and to generate diverse biobanks^1,5–9^, they are now also emerging as a translationally useful platform for mirroring clinical responses to therapy, such as in breast^10,11^, melanoma^12^, lung^13^, or gastrointestinal cancers^14–16^. However, while options for tumor organoid models exist, these organoids have not yet been used in the clinical setting in real time to aid with interpretation of treatment responses, particularly for central nervous system (CNS) tumors. Rather, they have been predominantly used for retrospective studies without direct application to temporally-matched patient treatments^10,11,14,17–19^. Therefore, whether tumor organoids can be generated sufficiently quickly and applied in real time to mirror clinical activity in patients and even to predict treatment responses for personalized medicine remains uncertain.

Glioblastoma (GBM) is the most common primary malignant brain tumor in adults and remains impervious to both standard of care and novel therapeutic methods^20^. While standard of care includes maximal surgical resection, radiation, and temozolomide chemotherapy, the survival of patients with primary GBM remains under two years^20^. For patients with recurrent GBM (rGBM), the survival time remains less than one year^20^, underscoring the immense unmet need for more effective therapeutic interventions. The extensive inter- and intra-tumoral heterogeneity of GBM not only drives therapeutic resistance, but also highlights the necessity for more high-fidelity experimental models beyond traditional monolayer cell cultures, which are limited in their ability to preserve cellular components and genetic profiles of parental tumors^3,19^. Additionally, traditional tumor modeling strategies, including xenografts and cancer cell lines, require a prolonged generation time that exceeds clinically relevant time windows. To overcome these obstacles, we previously developed a protocol for rapid generation of patient-derived glioblastoma organoids (GBOs) from fresh tumor specimens without single-cell dissociation, preserving key architectural elements of the original patient tumor that conventional two-dimensional and sphere culture methods lack^5,6^. More importantly, GBOs maintain key histological features, intra- and inter-tumoral heterogeneity, cell type and cellular state heterogeneity, and genetic mutation profiles of parent tumors as well as the tumor microenvironment, providing a high-fidelity model for clinical studies^5^. GBOs can also be biobanked and efficiently grafted into the adult mouse brain for xenograft studies^5,21^. Since our protocol can robustly generate GBOs within 2-3 weeks after surgery, it raises the possibility for real-time parallel assays with therapeutics given to patients to gain prospective insights into patient responses.

As a novel therapeutic modality, CAR-T cell therapies have been quite effective in hematologic cancers^22^, but have experienced limited success in solid tumors^23–25^, likely due to an immunosuppressive microenvironment and tumor antigen heterogeneity. To combat the cellular heterogeneity that contributes to disease recurrence in GBM, we have developed a bivalent CAR targeting both epidermal growth factor receptor (EGFR^26^) and interleukin-13 receptor alpha 2 (IL13Rα2^27,28^), which is currently in a phase 1 clinical trial in rGBM (NCT05168423)^29^. In this trial, patients first undergo a surgical debulking of the rGBM tumor, at which time an Ommaya reservoir is placed. Upon recovery from this surgery, patients are treated with a single intrathecal dose of EGFR-IL13Rα2 CAR-T cells. A recent interim analysis of this study demonstrated feasibility and safety of this therapy in the first 6 patients treated at doses of 1 × 10^7^ cells (*n* = 3) and 2.5 × 10^7^ cells (*n* = 3)^29^. In addition, correlative data demonstrated robust expansion of CAR-T cells in the cerebrospinal fluid (CSF) in all 6 patients, accompanied by inflammatory cytokine release. An early efficacy signal was also observed, with all 6 patients experiencing varying degrees of tumor regression as suggested by MR imaging within the first month following treatment. However, responses were not durable in all patients, and radiographic pseudoprogression was observed in a subset^29^, underscoring the emerging challenges associated with predicting clinical benefit from this novel therapy.

Determination of meaningful clinical efficacy for GBM treatment, in particular for CAR-T cells, may take months to manifest. Given this challenge, especially considering the short survival time associated with rGBM, real-time analysis of patient-derived GBOs could help to provide an early assessment of treatment efficacy and guide the next steps in patient management. Here we generated GBOs from the first six patients in the trial^29^ on the same timeline as production of the patients’ CAR-T cell products, all completed in the timeframe between surgical debulking of the tumor and treatment with the CAR-T cell product. These GBOs enabled simultaneous *ex vivo* testing of autologous patient CAR-T cells in parallel with patient treatment, providing crucial insights into therapeutic potency and patient responses. To the best of our knowledge, this is the first clinical trial with a unique design to perform patient-matched organoid correlative studies in real time with patient treatment. The inclusion and expansion of correlative patient-matched experiments are invaluable to the understanding of clinical efficacy and biological mechanisms behind these developing therapies.

## RESULTS

### Retention of CAR target tumor antigens in GBOs generated from clinical trial patient tissues

GBM tumor specimens were obtained from patients with rGBM enrolled in a phase I clinical trial (NCT05168423)^29^ under informed consent. In this trial, tumor tissue was collected from rGBM patients during surgical resection and Ommaya placement 2-4 weeks prior to investigational treatment with CAR-T cells (**Figure 1A**). GBOs were generated from fresh tumor tissue via micro-dissection and cultured in chemically-defined media^5,6^. The first six patients in the trial were examined, including patients 1-3 (dose level one, 1 × 10^7^ cells *in vivo*) and patients 4-6 (dose level two, 2.5 × 10^7^ cells *in vivo*) (**Table S1**). All patient tumor samples formed GBOs successfully within 2-3 weeks of tissue acquisition (**Figure S1A**). The experiments using GBOs in the current study were all performed concurrently with intrathecal autologous CAR-T cell infusions in the clinical trial (**Figure 1A**).

**Figure 1.**
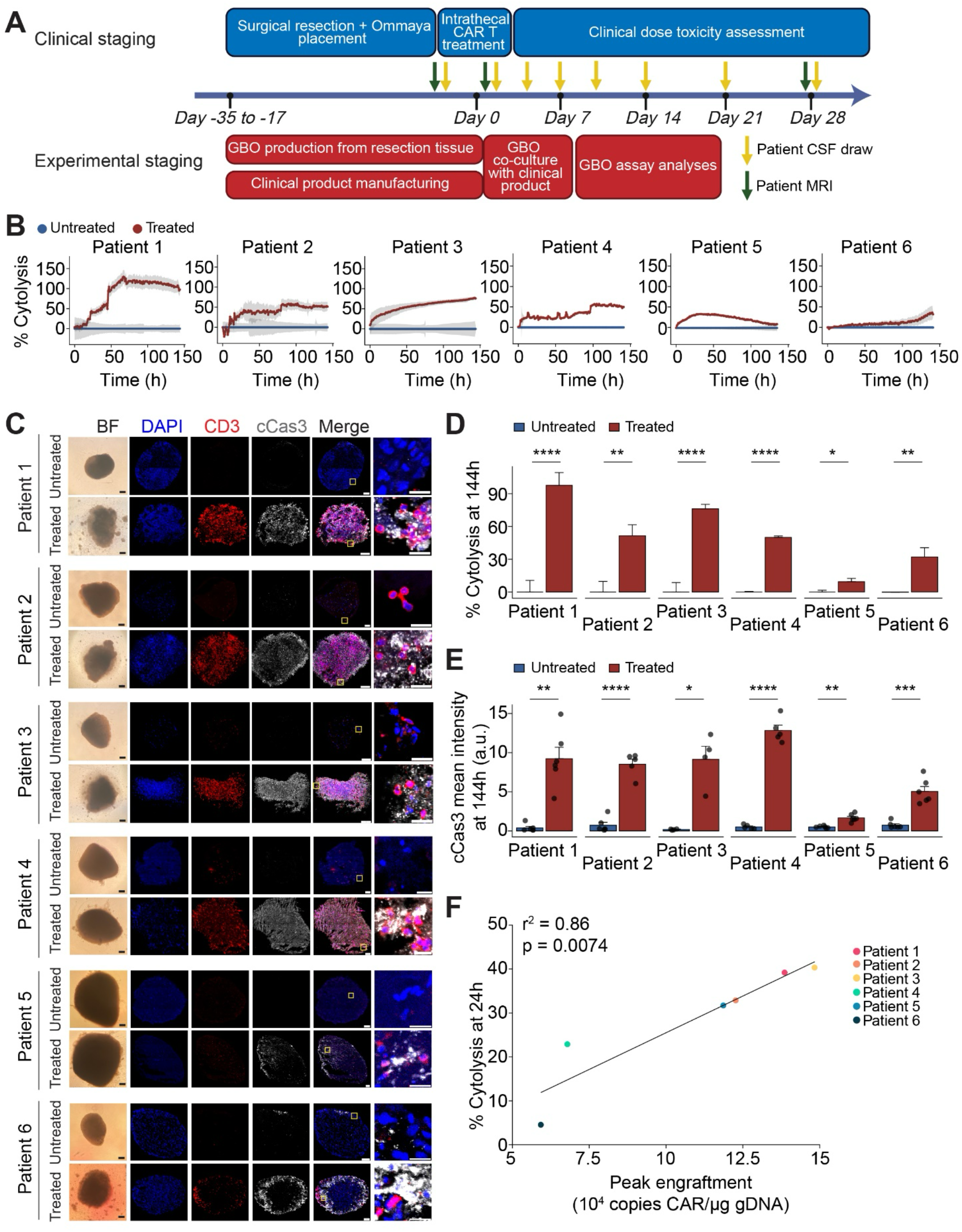
GBM organoid co-culture with patient-matched CAR-T cells revealed significant tumor cell cytolysis that correlated with clinical CAR-T cell engraftment. (**A**) Timeline scheme for correlative GBO assays performed in parallel to patient treatment in the clinical trial NCT05168423. (**B**) Time-course analysis of GBO cytolysis with co-culture of autologous CAR-T cells over 144 hours with the Axion Maestro Cellular Impedance platform, showing reduced impedance of GBOs by patient CAR-T cells (Treated) when compared to untreated GBOs (Control). Values represent mean ± S.E.M. (for all patients, *n* ≥ 6 GBOs for the control condition and n ≥ 12 GBOs for the treatment condition). (**C**) Sample images of bright field (BF) and immunofluorescence for DAPI, CD3, and cleaved-caspase 3 (cCas3) in treated and untreated GBOs, demonstrating both CD3^+^ T cell infiltration and cCas3^+^ tumor cell apoptosis in treated GBOs. Scale bars, 100 μm. Yellow squares denote inset merge images from respective GBOs (last panel). Scale bars, 20 μm. (**D**) Summary of endpoint cytolysis levels comparing autologous CAR-T cell treated to untreated GBOs. Same dataset as in (**B**). Values represent mean ± S.E.M. (\**p* < 0.05, \*\**p* < 0.01; \*\*\*\**p* < 0.0001; multiple Welch’s *t*-tests). (**E**) Quantification of cCas3 immunostaining in autologous CAR-T cell treated and untreated GBOs at 6 days post co-culture, confirming that values seen in (**B**) and (**D**) were due to apoptotic cell death. Values represent mean ± S.E.M. (n = 4-6 GBOs per condition; \**p* < 0.05, \*\**p* < 0.01; \*\*\**p* < 0.001; \*\*\*\**p* < 0.0001; multiple Welch’s *t*-tests). (**F**) Correlation plot of average cytolysis values at 24 hours in the treated GBOs compared to peak CAR-T cell engraftment levels in the corresponding patients. Average levels of CAR-T cells were detected in CSF post-CAR injections using qPCR (copies/μg gDNA). Strong positive Pearson correlation suggests that short-term activity in GBOs reflects the magnitude of CAR-T cell engraftment and/or proliferation in patients. Cytolysis values were from (**B**). Line was generated with a linear model fit, and values represent mean ± S.E.M. See also **Figures S1, S2**, and **S3**, and **Table S1**.

The targeted EGFR epitope (806) is expected on the surface of tumor cells in 50-60% of GBM patients, whereas IL13Rα2 is expressed in tumors in 50-75% of GBM patients^30–32^. Previous studies have shown that patient-derived GBOs retain the histologic features of parental tumors, including antigen heterogeneity^5^. We first performed immunohistochemistry analyses to confirm the presence and retention of both targeted tumor-associated antigens EGFR and IL13Rα2 in GBOs. EGFR and IL13Rα2 immunostaining signal in all six patient GBOs at 21 days after generation was qualitatively comparable to those in sections of parental tumors at day 0 (**Figures S1B-G**), suggesting the preservation of CAR-T targeted antigens in GBOs from parental tumor tissue.

### Tumor cell cytolysis in GBOs co-cultured with patient-matched CAR-T cells and correlation with clinical CAR-T cell engraftment

To perform real-time evaluation of CAR-T cell treatment response *ex vivo*, we collected patient CAR-T-EGFR-IL13Rα2 cells from the autologous infusion products obtained during quality control assessment (**Figure S1H**). Therapeutic testing was performed on at least 24 GBOs per patient from our phase I clinical trial protocol (NCT05168423)^29^. Using a rough effector to target (E:T) ratio of 1:10, autologous CAR-T cells and GBOs were co-cultured for 6 days within the Axion Maestro Cellular Impedance platform. The impedance assays measure the cytotoxic effects of CAR-T cells on GBOs by monitoring changes in electrical impedance as GBOs adhere to electrodes within the cell culture plates. Tumor cell killing was quantified using control impedance values of GBOs without treatment and full lysis impedance values over the same time points for each patient sample assay^33,34^. The end point was chosen based on the disruption of GBOs observed during co-culture. Each set of matched patient-derived GBOs and autologous CAR-T cells demonstrated significant tumor cytolysis over time as assessed by changes in electrical impedance (**Figures 1B, D**). Tumor cell killing was orthogonally confirmed by cleaved caspase 3 (cCas3) immunostaining in GBOs after 6 days of co-culture (**Figures 1C, E**).

Due to a tight clinical timeline combined with limited surgical tissue acquired from rGBM patients, we prioritized our experiments with the limited number of GBOs available. For a set of control experiments, we collected samples from an off-trial patient (UP-11273; removed from the trial due to patient performance status not being acceptable for CAR-T cell infusion) who also had autologous CAR-T cells generated (**Table S1**). We performed co-culture experiments using GBOs from this off-trial patient with autologous non-modified (untransduced, UTD) T cells, autologous CAR-T cells targeting CD19, and autologous bivalent CAR-T cells. We observed significant cytolysis induced by on-target CAR-T cells, but not CD19 CAR-T cells or UTD T cells, indicated by impedance killing assays as well as immunostaining for cCas3 in GBOs (**Figures S2A-C**).

Additional analysis of cCas3 immunostaining was performed at an earlier, 3-day timepoint to demonstrate continued and increasing anti-tumor activity over the total 6-day assay (**Figures S3A-B**). To further specify the targeted killing of tumor cells, we performed immunostaining of T cell marker CD3 and GBM tumor marker Nestin^35^. We found that CAR-T cells eradicated Nestin^+^ tumor cells, particularly at the edge of GBOs (**Figure S3C**).

To assess the degree to which *ex vivo* responses of GBOs matched patient responses *in vivo*, we considered the degree of engraftment of CAR-T-EGFR-IL13Rα2 cells in patient CSF as previously measured by quantification of CAR copies per microgram of genomic DNA^29^. Importantly, the magnitude of cytolysis at the 24-hour timepoint in impedance assays directly correlated with peak CSF engraftment in patients post-CAR-T cell administration^29^ for these six patients (**Figure 1F**). The matched 24-hour post-treatment magnetic resonance (MR) imaging revealed marked decreases in the intensities of contrast enhancement across all six patients^29^, which is suggestive, but not definitive, evidence for tumor killing. Our *ex vivo* GBO results therefore provide direct evidence for the efficacy of the CAR-T cell treatment in tumor killing.

### Activation and cytotoxic activity of patient-matched CAR-T cells in GBO co-cultures

Before use in co-culture assays, patient CAR-T cell products were analyzed for bivalent CAR expression after manufacturing (**Figure 2A**). Transduction efficiency was measured to ensure that patient products met the minimum release criteria of 2% CAR^+^ before administration, and we found robust expression of both individual CAR constructs in all six patient products (**Figure 2A**). In corroboration with CAR-T cell-mediated tumor killing, we conducted additional immunohistochemical staining to examine T cell penetration and expansion within treated GBOs. CD3 immunostaining of co-cultured GBOs showed the extent of T cell infiltration into GBOs after 6 days in culture (**Figure 2B**). After 6 days in co-culture, a large proportion of CD3^+^ cells in GBOs (>20% for all six patients) was positive for the proliferation marker Ki-67, suggesting continued activation of CAR-T cells across patients (**Figure 2C**). We verified that the proportion of Ki-67^+^ T cells was significantly higher than paired UTD T cells using off-trial GBO UP-11273 (**Figure S2D**). In line with persistent proliferation marker expression, CD3^+^ cells within the GBOs were observed to express the primary effector protease Granzyme B, which is secreted by activated T cells to mediate apoptosis in target cells^36^, and CD69, which is an early activation marker for T cells^37^ (**Figures S3D-E**). Collectively, in addition to tumor cytolysis, these results indicate activation and cytotoxicity displayed by patient CAR-T cells in co-culture with matched GBOs.

**Figure 2.**
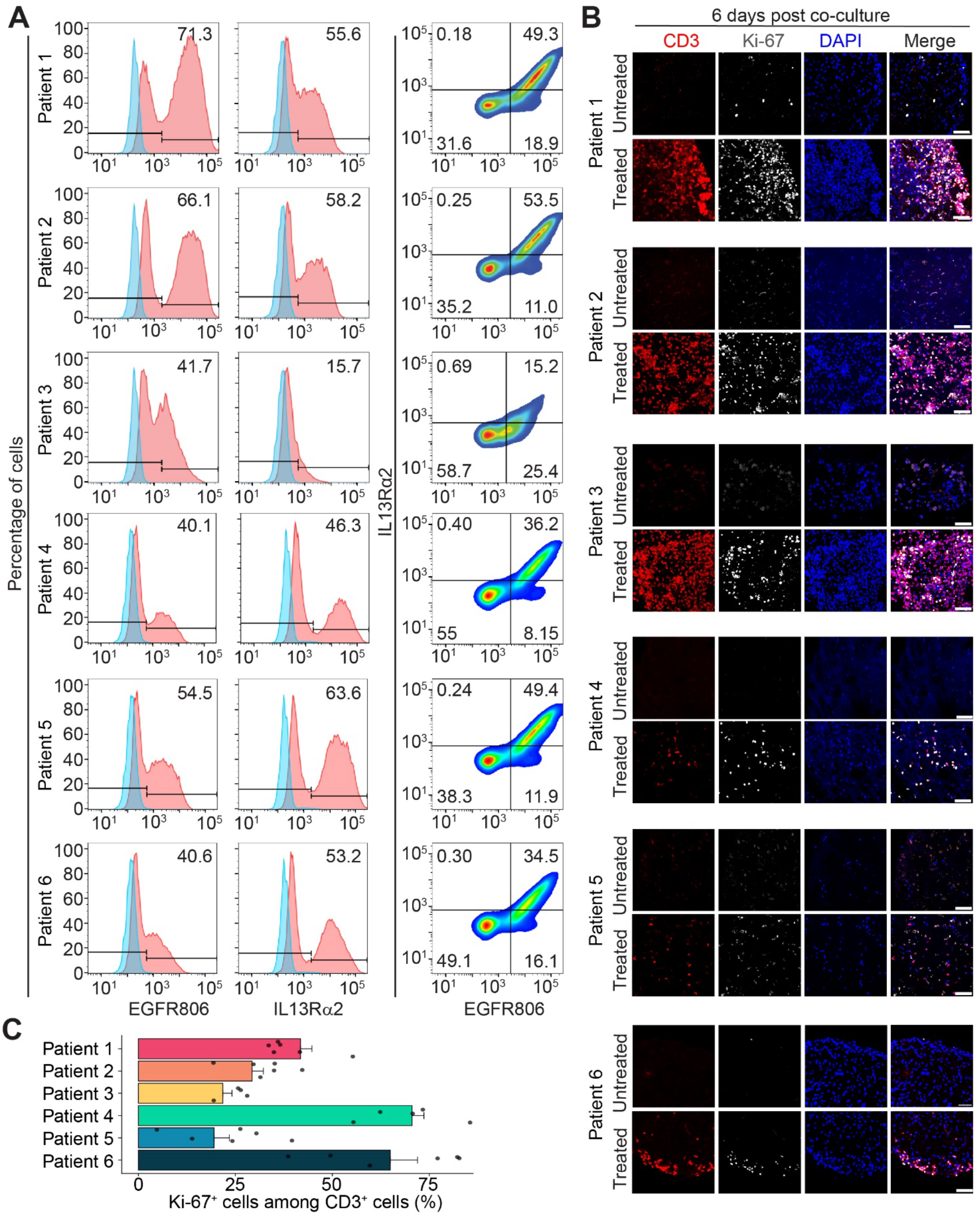
GBM organoid co-culture induced significant activation marker expression in patient-matched CAR-T cells. (**A**) Flow cytometric detection of patient T cell expression of each CAR from the bivalent construct (**Figure S1H**). Histograms depict EGFR CAR (left column) or IL13Rα2 CAR (center column) expression detected for each patient T cell product. Far right column shows plots to quantify dual CAR expression. (**B**) Sample immunofluorescence images for CD3, and Ki-67, and DAPI in treated and untreated GBOs at 6 days post co-culture, showing infiltration and actively proliferating CAR-T-EGFR-IL13rα2 cells. Scale bars, 50 μm. (**C**) Quantification of Ki-67 immunostaining in CAR-T-EGFR-IL13rα2 cells after 6 days of co-culturing with GBOs. Values represent mean ± S.E.M. (*n* = 5-6 GBOs per condition). See also **Figure S1**.

### Reduction of target antigens in post-treatment GBOs

Due to restricted access to patient tissue immediately after CAR-T cell treatment in the clinical trial, no information could be obtained regarding the status of target antigen presence on tumors in patients^29^. We therefore collected GBOs after 6 days in co-culture with matched patient CAR-T cells and analyzed the expression levels of target antigens in GBOs. Immunohistochemistry showed that levels of both EGFRwt (wild-type) and IL13Rα2 appeared to be reduced in GBOs treated with autologous CAR-T cell product (**Figures S4A-B**). To quantitatively evaluate changes more specifically in single or double antigen-positive cell populations, GBOs cultured alone or with autologous CAR-T cells were dissociated for flow cytometry analyses. Compared to isotype control immunostained GBO cells, treatment with autologous CAR-T cells led to a significant reduction of double-positive cell populations and a significant increase of double-negative cell populations across GBOs from patients (**Figures 3A-B**), which likely contributed to the substantial tumor cell cytolysis observed in our impedance assays (**Figure 1B**). Interestingly, we observed increased single IL13Rα2-positive and decreased single EGFR806-positive cell populations following treatment for four out of five patients examined (**Figure 3B**), which may suggest EGFR as the dominant driver of the activity of this bivalent CAR-T for most patients. Nonetheless, the heterogeneity of the single antigen-positive dynamics between patients underscores the benefits of the bivalent CAR-T design. Additionally, immunohistochemistry of off-trial control patient UP-11273 GBOs after CAR-T cell treatment showed colocalization of target antigen and cCas3, indicating on-target killing of antigen-positive tumor cells (**Figure S4C**). Notably, target antigen positivity was reduced only in GBOs treated with on-target CAR-T cells compared to CD19-targeting CAR-T cells or UTD T cells (**Figure S4D**). These results therefore suggest that the parallel organoid paradigm can provide direct measurement of and additional insights into the efficacy of CAR-T cells on rGBM.

**Figure 3.**
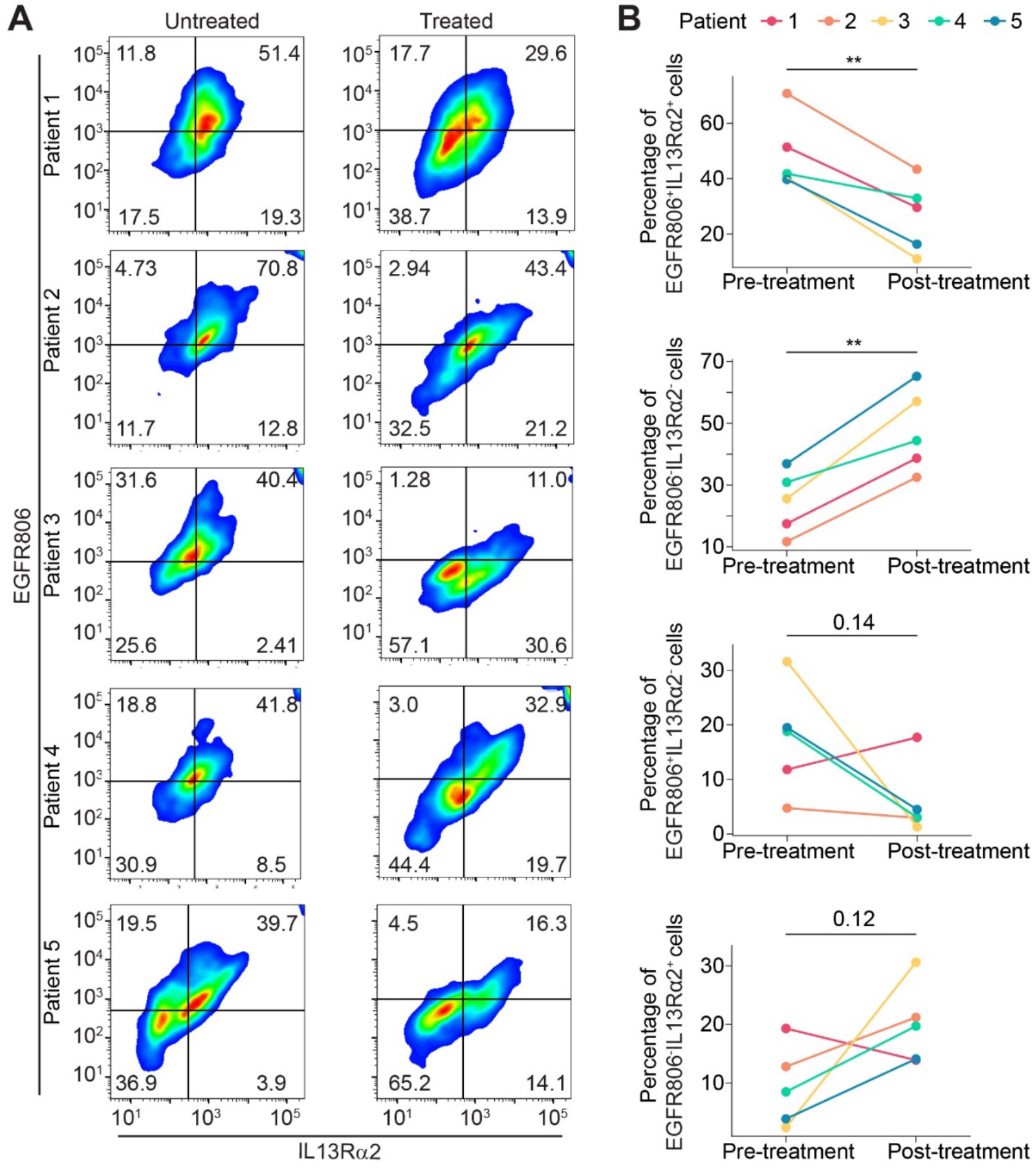
GBM organoid co-culture with patient-matched CAR-T cells exhibited reduction in target antigen expression. (**A**) Flow cytometry quantification of single or double target antigen-positive cells in dissociated GBO cells without (top) and with (bottom) co-culture treatment of CAR-T cells. Patient GBO flow data reveals dramatic reductions in double target positive cells after CAR-T cell treatment across all six patient GBO cohorts (top right quadrant) compared to single target positive EGFR^+^ tumor cells (top left quadrant) and IL13rα2^+^ tumor cells (bottom right quadrant). (*n* = 4 GBOs for each of patients 1-5; we did not have sufficient GBOs from patient 6 for this assay). (**B**) Quantification of antigen expression across GBOs before and after 6 days of co-culture with CAR-T-EGFR-IL13rα2 cells (*n* = 5 patients; \*\**p* < 0.01, paired Welch’s t-tests). See also **Figure S4**.

### Parallel time course of cytokine levels in CAR-T cell-GBO co-culture media and patient CSF

To further validate and quantify the cytotoxic activity by CAR-T cells, we collected and refreshed CAR-T cell-GBO co-culture media every day over 6 days and performed ELISAs on the co-culture conditioned media to detect IFNγ, TNFα, and IL-2 as metrics of T cell activation^38^. All cytokine values demonstrated a strong and continuous release by the activated T cells, with peaks occurring at days 1 or 2 (**Figures 4A-C**). In contrast, no cytokines were detectable in the blank control media or in media cultured with GBOs alone (**Figure S4E**). In addition, in GBOs from the off-trial control patient UP-11273, cytokines were minimally detectable in media from GBOs co-cultured with CD19 targeting CAR-T cells or UTD T cells, or media from T cells alone, compared to GBOs co-cultured with on-target CAR-T cells (**Figure S2E**).

**Figure 4.**
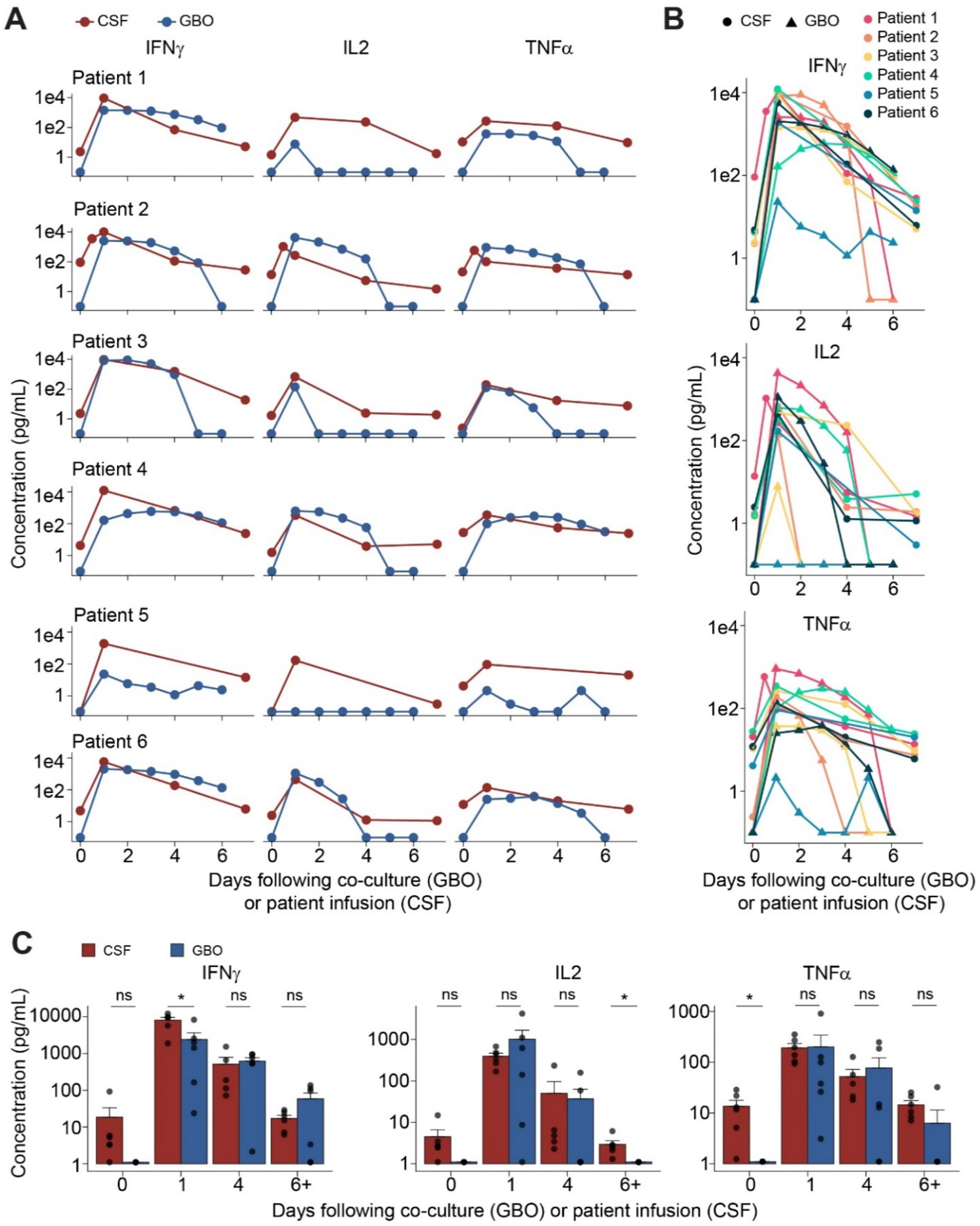
Patient-matched GBO media and patient CSF sampling exhibited commensurate trends in critical cytokines TNFα, IL-2, and IFNγ after CAR-T cell treatment. (**A**) Independent patient CSF and matched GBO cytokine data for IFNγ, IL-2, and TNFα levels showing peak release within the first four days post-treatment before returning to the baseline. (**B**) Combined trendlines for all patients and matched GBOs for each cytokine. (**C**) Comparison of patient CSF and matched GBO assay cytokine quantifications compared at each collection time point. For the 6+ day condition, Day 6 GBO concentrations were compared to Day 7 CSF concentrations. Values represent mean ± S.E.M. (*n* = 6 patients; ns: *p* > 0.05; \**p* < 0.05, multiple Welch’s *t*-tests with *p*-value adjustment by Bonferroni’s method). See also **Figures S1 and S4**.

Cytokine levels in GBO media were quantified and compared to paired patient CSF levels^29^. Across matching time points, TNFα, IL-2, and IFNγ trends in the GBO conditioned media were all commensurate with levels of these cytokines present in patient CSF (**Figures 4A-C**). Direct comparisons of GBO cytokines with matched patient CSF concentrations did not reveal statistically significant differences through 4 days of co-culture for all three cytokines (**Figure 4C**). Levels of cytokines within GBO assays demonstrated similar temporal dynamics compared to *in vivo* kinetics of CAR-T cell activation observed in patients in the clinical trial (**Figure 4B**).

## DISCUSSION

Our previous detailed characterizations have shown that patient-derived GBOs *ex vivo* resemble many features of patient GBM *in vivo*, including intertumoral heterogeneity, cell state diversity, landscape of the transcriptome and genomic mutations, and importantly, the tumor microenvironment^5^. Here we demonstrate proof-of-principle that GBOs can be rapidly generated from resected tissue for analyses even before patients receive matched treatment, providing real-time avatars to help interpret patient responses to novel therapies. We designed a novel paradigm of a clinical trial with treatment with autologous CAR-T cells in patients and in GBOs derived from the same patients *ex vivo*. Our study showed exceptional correlation of various parameters, including the degree of temporally matched GBO cytolysis *ex vivo* and CAR-T cell engraftment *in vivo*, and the time course of cytokine release *ex vivo* and in patient CSF. More importantly, our real-time analysis of GBOs *ex vivo* provides additional and critical insights into patient treatment responses. For example, our finding of direct and effective killing of tumor cells in GBOs by CAR-T cell treatment mitigates concerns about the occurrence of a “pseudo-response” in MR imaging^29,39,40^, increasing confidence about treatment efficacy. We also confirmed decreased levels of target antigens after CAR-T cell treatment, which is information not available clinically due to inaccessibility. As *in vivo* efficacy in patients takes months to manifest, such parallel analysis provides critical information to optimize ongoing and future clinical trials. Although confirmation of these exploratory, hypothesis-generating findings is needed with a larger sample size, our study suggests the potential for a broad application of our clinical trial design with simultaneous treatment of patients and patient-derived tumor organoids with the same clinical product for real-time monitoring of treatment responses and the generation of additional insights.

Glioblastoma remains a devastating disease characterized by a high rate of treatment resistance and recurrence^20^. Immunotherapy, particularly CAR-T cell therapy targeting solid tumors like glioblastoma, has encountered significant challenges due to tissue heterogeneity and a complex and immunosuppressive tumor microenvironment^24,25,41^. Our ability to overcome these challenges has been limited by a lack of experimental models that faithfully replicate the *in vivo* microenvironment. Building upon the observed correlation between treatment-induced responses of GBOs *ex vivo* and patients *in vivo*, our study suggests the potential of GBOs as a model to address these limitations. For example, the maintenance of antigen heterogeneity is a hallmark of GBM that was also observed within GBOs through the *ex vivo* presence of both antigen-positive and -negative tumor cells. Discovery of the mechanisms regulating tumor escape and subsequent recurrence following cellular immunotherapy is still greatly needed to inform future CAR-T cell therapies. By exploring single-versus double-target antigen-positive tumor cells after treatment, GBOs may thus provide such a model to determine tumor escape and recurrence mechanisms. These studies may validate the use of multivalent CAR-T cells and lead to additional approaches to further optimize the anti-GBM response *in vivo*. The elevated CSF cytokine levels of IFNγ, IL-2, and TNFα immediately after delivery support *in vivo* CAR-T cell expansion and cytotoxic activity, but these levels subsided after one week in all six patients^29^. Importantly, patient-derived GBOs exhibited a similar tempo of cytokine release, and recollection of CAR-T cells following co-culture with matched patient GBOs may be utilized to investigate T cell activation and exhaustion dynamics in order to develop new strategies to enhance the duration of CAR-T cell effectiveness. For instance, in an accompanying manuscript, we applied multi-omic single cell profiling approaches to dissect the temporal dynamics of interactions among CAR-T cells, tumor cells and tumor microenvironment cells in the CAR-T-GBO co-culture, which identified complex cell-cell interactions and signaling over time, and revealed unexpected anti-tumor effects on antigen-negative cells that may have therapeutic implications. While it is possible that 2D cultures could exhibit similar findings of antigen depletion, cytokine activation, and tumor cytotoxicity following CAR-T cell treatment, generation of these cultures are generally not possible within this quick timeframe and they normally lack crucial components of the tumor microenvironment^42,43^.

Together, our proof-of-principle study with a novel clinical trial design showed preliminary success in correlating *ex vivo* GBO responses with *in vivo* patient responses. These data provide a foundation for the future application of GBOs as avatars for testing treatment response in real time to stratify patients for clinical trials and to prioritize potential personalized treatment options. For example, it may be possible in the future to select an optimal CAR-T cell therapy from a portfolio of options or select from several candidate drugs. Given the short survival period after diagnosis for GBM and rGBM patients, such an approach could be critical to move the needle in treating this devastating disease.

### Limitations of the study

First, the current study is limited in the small sample size of the number of paired treatments of patients and patient-derived GBOs. Our analysis started with the cohort of the first three patients in the trial and we confirmed our findings from the initial cohort with a second cohort of next three patients, and we report the combined data in this paper. As the current clinical trial is ongoing, further studies will help to further validate and potentially enhance the current findings. Second, due to comparatively sparse surgical samples from rGBM patients and limited time for GBO expansion to match the clinical treatment timeline, we are constrained by the number of GBOs available and could not perform all sets of experiments for all patients. Third, we are limited by what samples can be collected throughout the course of optimized clinical care for GBM patients. Therefore, some correlatives cannot be accessed or measured *in vivo* at certain time points.

## ACKNOWLEDGEMENTS

We would like to thank the patients who participated in this study and their families who made this work possible. We thank the Neurosurgery Clinical Research Division, the Translational and Correlative Sciences Laboratory and the Clinical Cell and Vaccine Production Facility at the University of Pennsylvania School of Medicine for their contributions to the clinical trial and associated preclinical efforts. We also thank the Cell and Developmental Biology Microscopy Core Facility at University of Pennsylvania and Axion Biosystems for equipment support. This work was funded by the Abramson Cancer Center Pilot Award to M.L., the Abramson Cancer Center Glioblastoma Translational Center of Excellence to D.M.O., the Templeton Family Initiative in Neuro-Oncology to D.M.O., the Maria and Gabriele Troiano Brain Cancer Immunotherapy Fund to D.M.O., National Institutes of Health grants R35NS116843 to H.S., R35NS097370 to G-L.M., and F31NS137664 to Y.S., the Dr. Miriam and Sheldon G. Adelson Medical Research Foundation to G-L.M., and Institute for Regenerative Medicine and Department of Neurosurgery at the University of Pennsylvania to H.S., D.M.O., S.B. and M.N.

## AUTHOR CONTRIBUTIONS

H.S., D.M.O., G-L.M., Z.A.B., and S.J.B. conceptualized the study. H.S., D.M.O., G-L.M., and Z.A.B. supervised the study. D.M.O., M.P.N., and E.L.P. collected tumor tissue. X.W. and Y.S. generated and cultured the GBOs used in the study. M.L. provided and characterized CAR-T cell product and performed cytolysis analyses. X.W., M.L., and Y.S. performed CAR-T cell co-culture and analyses. X.W., M.L., and Y.S. performed immunohistochemical analyses. N.L., A.D., D.Y.Z., B.S.U., G.P., and D.S. performed additional experiments and analyses. H.S., D.M.O., G-L.M., and Z.A.B. acquired funding for the study. M.L., X.W., Y.S., H.S., D.M.O., G-L.M., and Z.A.B. wrote and edited the manuscript with input from all authors.

## Methods

### Human Subjects

All experimentation using human tissues described herein was approved by the Institutional Review Board at the University of Pennsylvania. GBM organoids (GBOs) were established as described previously^5^. Tissue used for GBO establishment was obtained from patients enrolled in a phase I trial, “CART-EGFR-IL13Rα2 in EGFR Amplified Recurrent GBM,” NCT05168423. Tissue was obtained from in a total of seven patients (**Table S1**) with patient written consent, 26 to 30 days prior to CAR-T cell administration, at the time of Ommaya placement. One of these patients (UP-11273) was eventually taken off the trial due to the performance status not being acceptable for CAR-T treatment infusion.

### GBO Generation

Co-culture experiments were performed within 2-3 weeks after GBO establishment to ensure the retention of the tumor microenvironment in the organoids^5^. GBOs were generated and maintained as described previously^5,21^. In brief, freshly resected surgical rGBM tissue was placed in dissection medium (Hibernate A (Thermo Fisher), 1X GlutaMax (Thermo Fisher Scientific), and 1X Antibiotic-Antimycotic (Thermo Fisher Scientific)) prior to transport to the lab, where tissue was dissected into small (∼1 mm^3^) pieces. Dissected tissue pieces were washed in 1X Red Blood Cell lysis buffer (Thermo Fisher Scientific) and washed in DPBS prior to culture in a 6-well plate in GBO culture medium (50% Neurobasal (Thermo Fisher Scientific), 50% DMEM:F12 (Thermo Fisher Scientific), 1X NEAAs (Thermo Fisher Scientific), 1X GlutaMax (Thermo Fisher Scientific), 1X Penicillin-Streptomycin (Thermo Fisher Scientific), 1X B27 without vitamin A supplement (Thermo Fisher Scientific), 1X N2 supplement (Thermo Fisher Scientific), 1X 2-mercaptoethanol (Thermo Fisher Scientific), and 2.5 μg/ml human recombinant insulin (Sigma)). Tissue pieces were cultured as GBOs on an orbital shaker (110 rpm) in a 37°C, 5% CO_2_, and 85% humidity sterile incubator until further analysis.

### Tumor Cytolysis Assays

CytoView 96-well impedance plates (Axion Biosystems, Atlanta, GA) were prepared by coating with 0.1 mg/mL poly-D-lysine (Sigma Aldrich, Burlington, MA) then 20 µg/mL laminin (Sigma Aldrich) overnight at 37°C. GBOs were plated onto CytoView plates and monitored by the Maestro ZHT platform (Axion Biosystems) for 2 days to ensure adherence. For co-cultures, 1:10 patient T cells: tumor cells were added, and data were collected every minute for 6 days. The E:T ratio was generated using an estimation of tumor cells in an organoid^5^. Variability in total tumor cell numbers of a GBO was accounted for by the large number of biological replicates, with a minimum of 12 GBOs run per condition. Changes in impedance are reported as the resistive component of the complex impedance recordings within AxIS software (Axion Biosystems) and all data are corrected to the impedance at the time of addition of T cells. Percent cytolysis calculations use untreated and full lysis controls to determine target cell cytolysis at collected time points as follows:

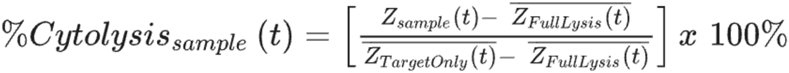

All cytolysis calculations were generated using untreated and full lysis control data within associated AxIS software. If the normalized impedance values outperformed the lysis controls, target cell killing values can exceed 100%.

### Clinical Vector Manufacturing

The bicistronic CAR-T-EGFR-IL13Rα2 vector encodes two CAR transgenes that recognize either the EGFR epitope 806^26^ or IL13Rα2^28^. Both transgenes are expressed from the same transcript using P2A, the 2A self-cleaving mechanism of porcine teschovirus (**Figure S1H**). The CAR sequences were cloned into a third-generation self-inactivating lentiviral expression vector containing the EF-1a promoter, a cPPT sequence, a Rev response element, and a woodchuck hepatitis virus posttranscriptional regulatory element (WPRE). Plasmid DNA was generated by Puresyn, Inc. The lentiviral vector was produced via transient transfection with four plasmids expressing the transgene, RSV-Rev, VSV-G and gag-pol in human embryonic kidney 293T cells at the University of Pennsylvania Center for Advanced Retinal and Ocular Therapeutics Vector Core.

### GBO and Tissue Immunohistochemistry

Serial tissue sections (20 μm thickness) of GBOs were sliced using a cryostat (Leica, Deer Park, IL), and melted onto charged slides (Thermo Fisher Scientific). Slides were dried at room temperature and stored at −20°C until ready for immunohistology. For immunofluorescence staining, the tissue sections were outlined with a hydrophobic pen (Vector Laboratories, Burlingame, CA) and washed with TBS containing 0.1% Tween-20 (v/v). Tissue sections were permeabilized and non-specific binding was blocked using a solution containing 10% donkey serum (v/v), 0.5% Triton X-100 (v/v), 1% BSA (w/v), 0.1% gelatin (w/v), and 22.52 mg/ml glycine in TBST for 1 hour at room temperature. The tissue sections were incubated with primary antibodies diluted in TBST with 5% donkey serum (v/v) and 0.1% Triton X-100 (v/v) overnight at 4°C. Primary antibodies were mouse monoclonal anti-CD3 (BioLegend, San Diego, CA), rat monoclonal anti-CD3 (Genetex, GTX11089-100), mouse monoclonal anti-Granzyme B (Novus Biologicals, MAB2906), rabbit monoclonal anti-Nestin (Abcam, Ab105389), rabbit polyclonal anti-cleaved caspase 3 (Cell Signaling Technology, 9661S), rabbit monoclonal anti-Ki67 (Abcam, ab16667), and rabbit polyclonal anti-CD69 (Proteintech, 10803-1-AP). After washing in TBST, the tissue sections were incubated with secondary antibodies and DAPI (Sigma Aldrich) diluted in TBST with 5% donkey serum (v/v) and 0.1% Triton X-100 (v/v) for 1.5 hours at room temperature. Secondary antibodies were donkey polyclonal anti-mouse IgG (Thermo Fisher Scientific) and donkey polyclonal anti-rabbit IgG (Thermo Fisher Scientific). After washing with TBST, sections were incubated with TrueBlack reagent (Biotium, Fremont, CA) diluted 1:20 in 70% ethanol for 1 minute to block autofluorescence due to lipofuscin and blood components. After washing with DPBS, slides were mounted in mounting solution (Vector Laboratories), cover slipped, and sealed with nail polish.

For detection of antigen levels in treated or untreated GBOs, the primary antibodies used were rabbit monoclonal anti-EGFR (Cell Signaling, 4267S) and goat polyclonal anti-IL13Rα2 (R&D Systems, AF146). Images were acquired on a Zeiss LSM 710 confocal microscope and analysis was performed using ImageJ Fiji 2.9.0.

### Flow Cytometry

To characterize tumor antigen presence additionally via flow cytometry, GBOs were harvested from co-culture assays and dissociated into single cell suspensions gently by incubation for 20 mins at room temperature in 0.5M EDTA, pH 8.0, solution (Invitrogen). The cells were then stained with live/dead exclusion marker DAPI solution (BD Pharmingen), followed by staining with the anti-EGFR806 Rabbit IgG (Absolute Antibody) and anti-IL13Rα2 Mouse IgG (R&D) antibodies at 4°C for 30 minutes. Cells were washed and stained for secondaries using Alexa Fluor Anti-Rabbit 488 and Alexa Fluor Anti-Mouse 647 before analysis on a 5-laser LSRFortessa flow cytometer and analyzed using FlowJo 10.10.0 software. Cells were gated on the live population, size, singlets by FSC, singlets by SSC before analysis for EGFR806 and IL13Rα2 presence.

Samples of patient CAR product were tested for both EGFR806 and IL13Rα2 scFv expression by staining with biotinylated EGFRvIII (AcroBiosystems) and fc-tagged IL13Rα2 (R&D) proteins. Binding to CAR was subsequently detected using streptavidin-conjugated FITC and anti-fc Alexa Fluor 647 (Biolegend). The cells were acquired on a 5-laser LSRFortessa flow cytometer and analyzed using FlowJo 10.10.0 software. Cells were gated on the live population, size, singlets and then CAR expression.

### Cytokine Quantification

For IFNγ, TNFα, and IL-2 detection in GBOs, the media from co-culture assays was collected and refreshed daily until co-culture day 6. Media of each condition (GBO alone, CAR-T cells alone, autologous CART-EGFR-IL13Rα2 cells with GBOs, or allogeneic CAR-T-EGFR-IL13Rα2 cells with GBOs) was pooled and measured by ELISA detection kits. All data were obtained with at least *n* = 2 technical replicates per condition.

Patient serum and CSF supernatant sampling was collected pre- and post-CAR-T cell treatment and cryopreserved at -80°C. Samples were thawed and analyzed for multiplex cytokine measurements using a custom 32-plex human cytokine panel designed for the trial by EMD Millipore Corporation. The panel included: EGF, FGF-2, Eotaxin, sIL-2Ra, G-CSF, GM-CSF, IFN-α2, IFN-γ, IL-10, IL-12P40, IL-12P70, IL-13, IL-15, IL-17A, IL-1RA, HGF, IL-1β, CXCL9/MIG, IL-2, IL-4, IL-5, IL-6, IL-7, IL-18, CXCL8/IL-8, CXCL10/IP-10, CCL2/MCP-1, CCL3/MIP-1α, CCL4/MIP-1β, RANTES, TNF-α, and VEGF. Samples were analyzed in duplicate according to the manufacturer’s instructions and compared against a six-point standard curve within internal standards. Data were acquired on a FlexMAP-3D system (Luminex) and analysis was performed on XPonent 4.3 software (Luminex). For any low out-of-range values, the baseline readings were approximated as done previously^22,29^. Measurement of some patient cytokines were reported in the recent study^29^.

### Quantification and Statistics

For GBO experiments, Welch’s *t*-tests were used to compare CAR-T-treated vs. untreated GBO conditions, and Pearson correlation was used to assess the relationship between the percent cytolysis in CAR-T-treated GBOs and CSF CAR-T cell engraftment in patients. T tests with adjustment (Bonferroni) for multiple comparisons were used to compare IFNγ concentration in GBO co-culture with CAR-T cells (**Figure S4**), and to compare GBO cytokine levels versus CSF concentrations at various days (**Figure 4C**). Quantification of a subset of immunohistochemistry data (Ki-67 and cCas3) was performed by semi-automated scripts developed in ImageJ/FIJI (see accompanying manuscript). No data were excluded from the analyses, and the correlative experiments were not randomized. The investigators were not blinded to allocation during experiments and outcome assessment. Data distribution was assumed to be normal, but this was not formally tested. *P* < 0.05 was considered significant. All statistical tests described were two-sided and performed using R version 4.1.2 (R Foundation for Statistical Computing, Vienna, Austria).

