## Supplementary Information for "Patient-derived glioblastoma organoids as real-time avatars for assessing responses to clinical CAR-T cell therapy"

**Inventory**

Figures S1-4

Table S1 legend

### SUPPLEMENTAL FIGURES

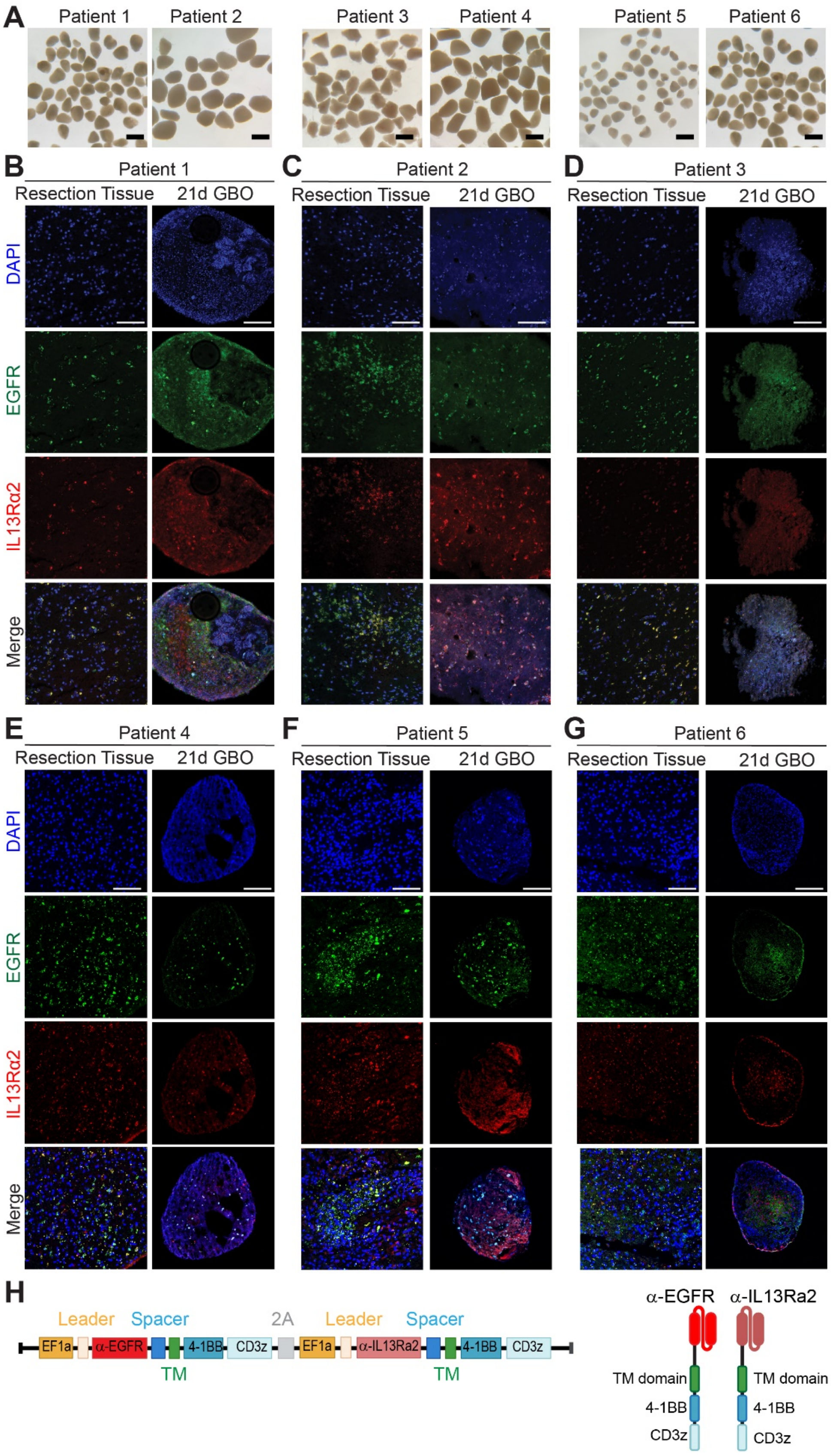

**Figure S1. Characterization of CAR target antigen expression in pre-infusion patient tumor tissue and matched patient-derived GBOs, related to Figure 1**.

**(A)** Sample brightfield microscopy images of GBOs generated from patients 1 through 6 at 21 days post tissue acquisition. Scale bars, 1 mm.

**(B-G)** Tumor tissue obtained at the time of Ommaya placement and matched GBOs at day 21 were immunostained for both CAR target antigens (EGFR in green, IL13Rα2 in red, and dual-positive cells in yellow). All 6 patients demonstrated expression of one or both targets throughout their tumor. All 6 patient tumor samples formed GBOs within 2-3 weeks of tissue acquisition and retained target antigen expression of EGFR and IL13Rα2 in *ex vivo* culture. See **Figure S4B** for quantifications. Scale bars, 200 μm.

**(H)** Schematic vector map of the bivalent CAR-T-EGFR-IL13Rα2 construct (left) that creates two parallel expressed CARs (right) within the cell after transduction. Antigen-binding domains shown are a-EGFR and a-IL13Ra2, which are single-chain variable fragments (scFv) molecules derived from monoclonal antibodies. Each scFv contains an intracellular signaling domain, consisting of a T cell activation domain derived from the CD3z chain of a T cell receptor as well as a 4-1BB (CD137) co-stimulatory domain. The constructs contain a dissolvable 2A linker sequence that allows the constructs to be expressed separately on the cell surface. Made using Biorender.com.

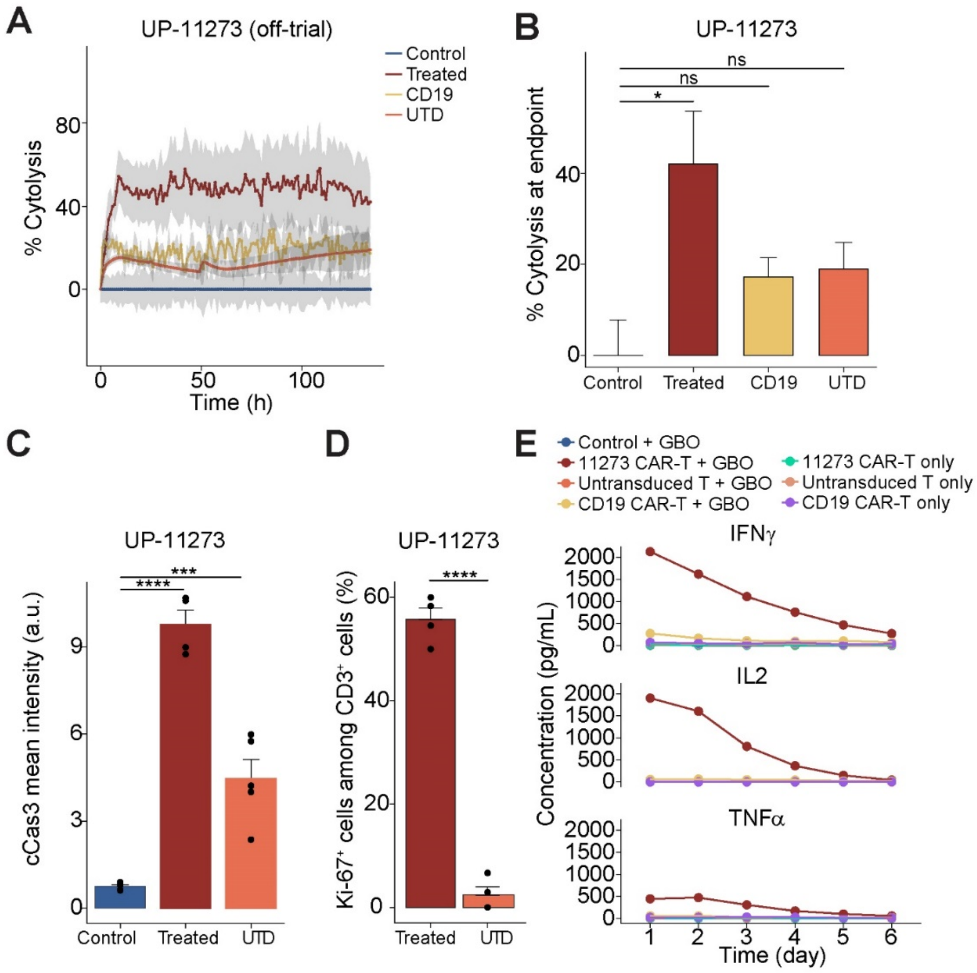


**Figure S2. Significant tumor cytolysis and T cell cytokine release in co-culture of off-trial UP-11273 GBOs with patient-matched on-target CAR-T cells, but not in co-culture with patient-matched off-target CD19 CAR-T cells or untransduced T cells, related to Figures 1 and 4.**

**(A**) Time-course analysis of GBO cytolysis assay of off-trial patient GBO UP-11273 in co-culture with autologous on-target EGFR-IL13Ra2 CAR-T cells, autologous off-target CD19 CAR-T cells, or autologous untransduced (UTD) T cells over 144 hours, showing specific cytolysis by patient on-target CAR-T cells, but not off-target CD19 CAR-T cells, UTD T cells, or untreated controls.

**(B)** Endpoint cytolysis levels at 144 hours of co-culture between GBOs and patient-matched on target bivalent CAR-T cells, off-target CD19 CAR-T cells, UTD T cells, or untreated GBO controls. Values represent mean ± S.E.M. (On-target CAR-T: $n$ = 13; CD19 CAR-T: $n$ = 8; UTD: $n$ = 6; Untreated: $n$ = 3 GBOs; ns: $p > 0.05$, *$p < 0.05$, multiple Welch's $t$-tests).

**(C)** Quantification of immunostaining performed on GBOs after co-culture with CAR-T-EGFR-IL13rα2 cells at the 6 day endpoint displays increased mean intensity of cCas, indicating increased apoptotic cell death due to CAR-T-EGFR-IL13rα2 compared to GBOs treated with matched-patient

UTD T cells or left untreated control. Values represent mean ± S.E.M. (*n* = 4-6 GBOs per condition; ***$p < 0.001$; ****$p < 0.0001$; multiple Welch's *t*-tests).

**(D)** Quantification of Ki-67 immunostaining in CAR-T-EGFR-IL13rα2 cells and patient-matched UTD T cells after 6 days in co-culture with GBOs. Values represent mean ± S.E.M. (*n* = 5-6 GBOs per condition; ****$p < 0.0001$; Welch's *t*-test).

**(E)** Cytokine levels from GBO co-culture media for IFNγ, IL-2, and TNFα over 144 hours, showing release of cytokines only in co-cultures containing CAR-T-EGFR-IL13rα2 cells and matched GBOs. No cytokine release was detected in cultures of T cells alone or in co-cultures containing GBOs with off-target CD19 CAR-T cells or UTD T cells.

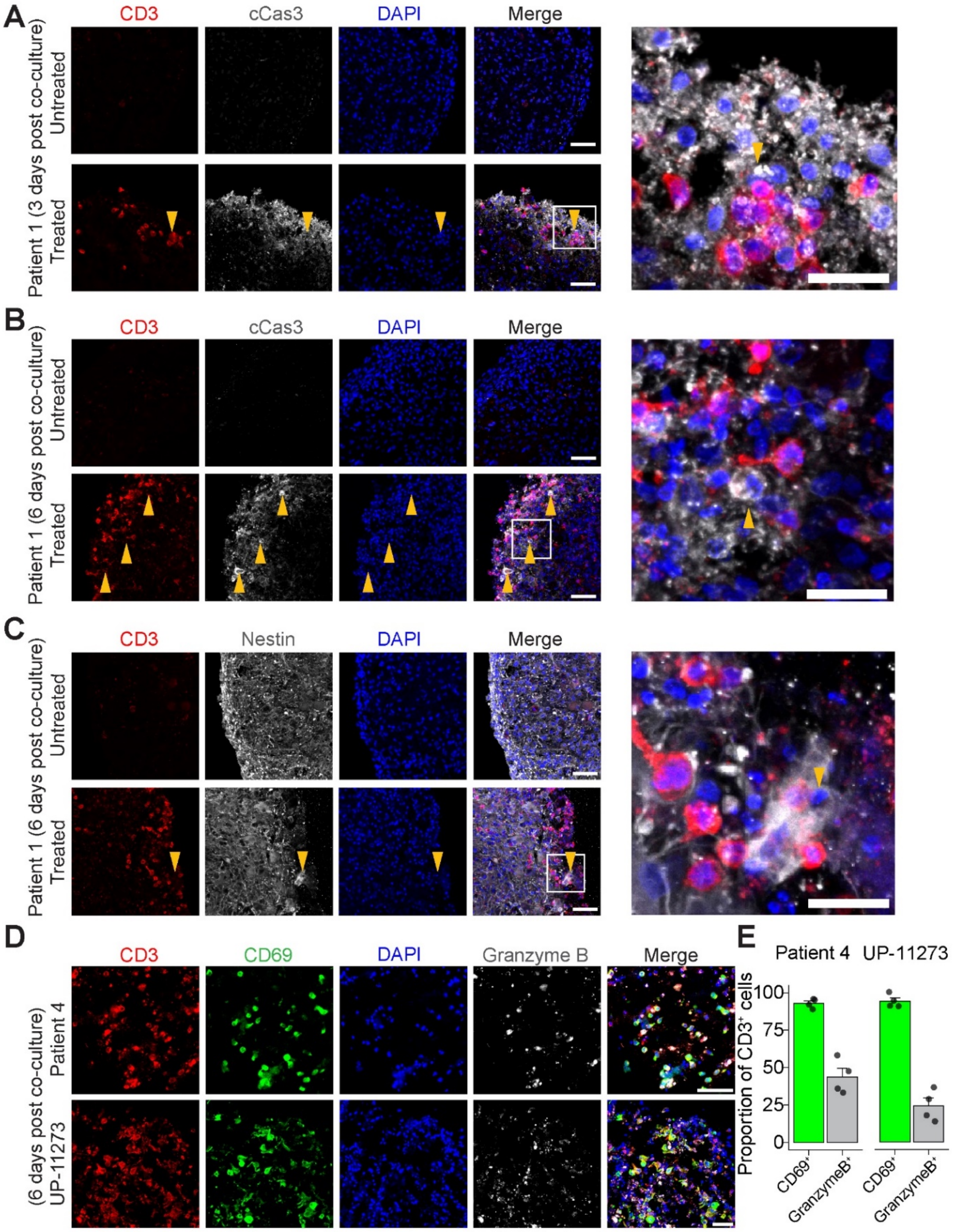


**Figure S3. Induction of robust tumor cytolysis and CAR-T cell activation in co-cultures of GBOs and patient-matched CAR-T cells, related to Figure 1.**

**(A-B)** Sample immunofluorescence images of CD3, cCas3, and DAPI in treated and untreated GBOs at 3 days **(A)** and 6 days **(B)** post co-culture with patient CAR-T cells. Orange arrowheads indicate cCas3$^+$ GBO cells surrounded by CD3$^+$ CAR-T cells, showing strong tumor cytolysis. Scale bars, 50 μm.

**(C)** Sample immunofluorescence images of CD3, NESTIN, and DAPI in treated and untreated GBOs at 6 days post co-culture with patient CAR-T cells. Orange arrowheads indicate the morphology change exhibited by NESTIN$^+$ tumor cells undergoing cytolysis. Note that CAR-T cells eradicated the NESTIN$^+$ cells along the GBO edge compared with that in the untreated organoids. Scale bars, 50 μm.

**(D)** Sample immunofluorescence images of CD3, CD69, Granzyme B and DAPI in treated and untreated GBOs at 3 days post co-culture, confirming presence and activation of T cells within GBOs. Patient 4 from the trial shown on top, and off-trial patient UP-11273 shown below. Scale bar, 50 μm.

**(E)** Quantification of CD69$^+$/CD3$^+$ or GranzymeB$^+$/CD3$^+$ T cells remaining within GBOs at 6 days post co-culture, indicating levels of activation among infiltrating T cells for Patient 4 and off-trial GBO UP-11273. Values represent mean ± S.E.M. ($n$ = 4 GBOs per condition).

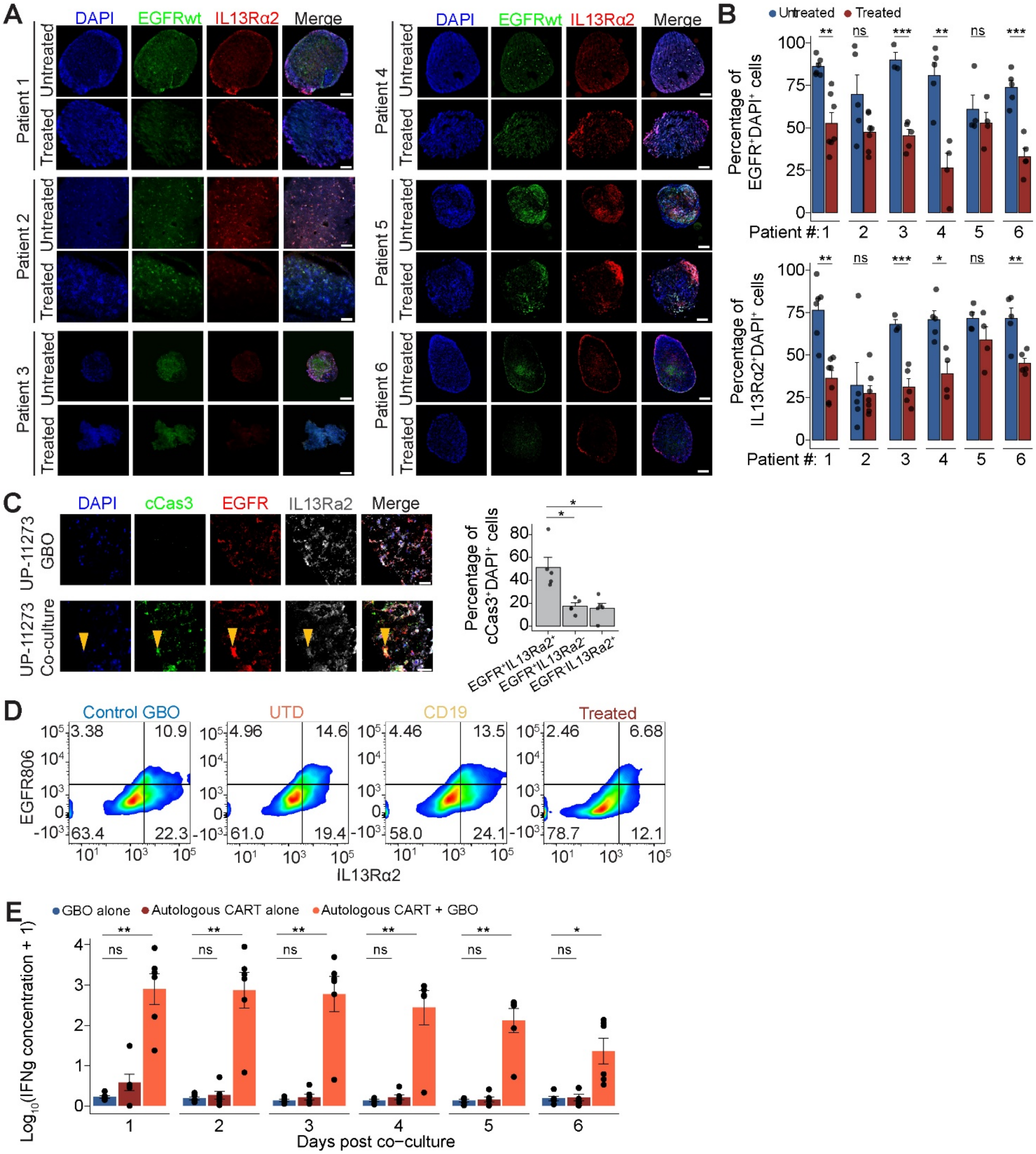


**Figure S4. CAR-T cell-GBO co-cultures display a decrease in tumor target antigen expression by 6 days and dynamic IFNγ release over time, related to Figures 3 and 4.**

**(A)** Sample immunofluorescence images for DAPI, wild-type EGFR, and IL13rα2 in treated and untreated GBOs after 6 days in co-culture, demonstrating qualitative reduction in target antigen expression in GBOs treated with CAR-T-EGFR-IL13rα2 cells in all 6 patients. Scale bars, 100 μm.

**(B)** Quantification of levels of EGFR and IL13rα2 immunofluorescence in treated and untreated GBOs. Values represent mean ± S.E.M. ($n$ = 4-8 GBOs per condition. ns: $p > 0.05$, *$p < 0.05$, **$p < 0.01$, ***$p < 0.001$; multiple Welch's $t$-tests).

**(C)** Sample immunofluorescence images for DAPI, cCas3, EGFR and IL13rα2 in treated and untreated GBOs of off-trial patient 11273 (left). Scale bars, 50 μm. Shown on the right is the quantification of levels of $EGFR^+$ and IL13rα$2^+$ in $cCas3^+$ cells, indicating that dying cells expressed one or both CAR antigens. Values represent mean ± S.E.M. ($n$ = 4-6 GBOs per condition; *$p < 0.05$; multiple Student's $t$-tests).

**(D)** Flow cytometry analysis of single or double target antigen-positive cells in dissociated GBO cells with and without co-culture of CAR-T cells. Target antigen expression only changed in GBO cells after treatment with on-target CAR-T-EGFR-IL13rα2 cells and did not change from baseline in either autologous CD19 CAR-T cells or UTD T cells co-cultures ($n$ = 4 GBOs per treatment).

**(E)** Daily measurements of IFNγ released in the culture media from GBOs alone (blue), autologous CAR-T-EGFR-IL13rα2 cells alone (red), or co-culture of autologous CAR-T-EGFR-IL13rα2 cells with GBOs (orange) for all six patients. Data are plotted as $\log_{10}(IFN\gamma + 1)$, where IFNγ is the concentration measured in pg/mL for each patient per condition ($n$ = 6 patients; ns: $p > 0.05$, ***$p < 0.001$, ***$p < 0.001$, multiple Welch's $t$-tests with $p$-value adjustment by Bonferroni's method).

### SUPPLEMENTARY TABLE

**Table S1. Patient information, related to Figure 1 (in Excel).**

**(A)** Patient basic information.

**(B)** Disease associated mutation panel.
